# Wilms Tumorigenesis in Human Kidney Organoids

**DOI:** 10.1101/2021.02.02.429313

**Authors:** Verena Waehle, Rosemarie Ungricht, Philipp S. Hoppe, Joerg Betschinger

**Affiliations:** Friedrich Miescher Institute for Biomedical Research, CH-4058 Basel; Faculty of Sciences, University of Basel, CH-4003 Basel; Novartis Institutes for BioMedical Research, Novartis Pharma AG, CH-4056 Basel

## Abstract

The loss or failure of cell differentiation is a hallmark of cancer, yet whether perturbation of differentiation is causal or consequential to malignant transformation is largely unclear. Wilms tumor is the most widespread kidney cancer in children. Here, we establish a model for Wilms tumorigenesis in human kidney organoids. We show that loss of the tumor suppressor *WT1* during organoid formation induces overgrowth of kidney progenitor cells at the expense of differentiating tubules. Functional and gene expression analyses demonstrate that absence of *WT1* halts progenitor cell progression at a pre-epithelialized cell state and recapitulates the transcriptional changes detected in a subgroup of Wilms tumor patients with ectopic myogenesis. By “transplanting” *WT1* mutant cells into wild-type kidney organoids, we find that their propagation requires an untransformed microenvironment. Genetic engineering of cancer lesions in human organoids therefore permits phenotypic modeling of tumor initiation and progression, and complements the current toolbox of pre-clinical Wilms tumor models.

## INTRODUCTION

Tumor initiation and progression is typically studied in genetically engineered mice (Kersten *et al*, 2017), whose relevance to human disease is limited by species- and strain-specific mechanisms, and the artificial induction of multiple oncogenes at the same time. Most human cancer models, in contrast, employ patients’ cells that are derived from tumor resections or biopsies, typically at a late stage of the disease, and that are grown as cell lines on tissue culture plastic in 2D, as tumor spheres in 3D, or as patient-derived xenografts (PDX) in mice (Hynds *et al*, 2018). Continuous growth and evolution of tumor cells in non-physiological environments imposes artificial selections (Ben-David *et al*, 2019) that can expose vulnerabilities irrelevant to the disease, which are thought to contribute to the poor clinical predictiveness of cancer models, in particular of tumor cell lines. To enhance clinical translatability, preclinical models are needed that more accurately represent the complexity of human cancers, such as tumor etiology, tumor heterogeneity and tumor environment.

Patient tumor-derived organoids preserve tumor heterogeneity, stages of tumor progression and drug responses (Clevers & Tuveson, 2019). Conversely, engineering of cancer lesions into wild-type organoids induces growth factor-independent growth and predisposes to tumor formation upon xenotransplantation (Kawasaki *et al*, 2020; Fumagalli *et al*, 2017; Bian *et al*, 2018; Ogawa *et al*, 2018; Matano *et al*, 2015; Drost *et al*, 2015). Acquisition of disease traits is, in fact, evident in engineered organoids prior to transplantation: Introduction of colorectal cancer mutations into intestinal organoids, and of brain cancer lesions into induced pluripotent stem cell (iPSC)-derived cortical organoids were shown to promote transcriptional signatures that resemble premalignant adenoma (Matano *et al*, 2015), and glioblastoma and primitive neuroectodermal tumors (Ogawa *et al*, 2018; Bian *et al*, 2018), respectively. Similarly, addition of TGFβ to intestinal organoids derived from adenomas directs progression into a mesenchymal colorectal cancer phenotype (Fessler *et al*, 2016). Organoids may therefore be suitable to model tumor initiation and progression *in vitro*. Here, we test this possibility by introducing Wilms tumor patient lesions into iPSC-derived human kidney organoids.

Wilms tumor is the most common kidney cancer in childhood and accounts for about 7% of all pediatric cancers (Treger *et al*, 2019). Stalled nephrogenesis is thought to be the major cause of disease. This is supported by the transcriptional similarity of Wilms tumors with fetal cell types and by the function of several Wilms tumor oncogenes and tumor suppressors in normal kidney development. The diversity of genes that are mutated in Wilms tumor patients is larger than in other childhood tumors, and includes homozygous inactivation of the tumor suppressor Wilms tumor 1 (*WT1*), biallelic expression of insulin-like growth factor 2 (*IGF2*), neomorphic mutations in genes encoding the kidney transcription factors (TFs) SIX1 and SIX2, activation of the TF β-Catenin and disruption of miRNA biogenesis. The reason for this diversity is unclear. Some mutations, likely those that occur infrequently, may be associated with later stages of tumor development (Treger *et al*, 2019). However, gene expression analysis has suggested that Wilms tumorigenesis initiates in distinct cell types of origin and is influenced not only by which genes or pathways are mutated, but also by the developmental context in which genetic lesions occur (Gadd *et al*, 2012). Nephrogenesis may therefore be particularly vulnerable to transformation.

Here, we exploited human kidney organoids and showed that genetic knockout (KO) of *WT1* induces overgrowth of nephron progenitor cells (NPCs) at the expense of tubular and glomerular differentiation. Further characterization revealed progression into an organoid state that transcriptionally and phenotypically resembles a subtype of Wilms tumor patients, as well as arrest of NPC differentiation at a pre-epithelialized cell state. Our study therefore establishes modeling of Wilms tumor initiation and progression in human kidney organoids, and defines transcriptional and phenotypic hallmarks of cellular transformation in the absence of *WT1*.

## RESULTS

### *WT1* deletion inhibits NPC epithelialization and differentiation, and induces organoid hyperplasia

We generated kidney organoids using an adaptation of a two-step differentiation protocol (Morizane *et al*, 2015) (Ungricht et al., unpublished) that steers pluripotent iPSCs via intermediate mesoderm into SIX2-expressing NPCs within 9 days (d) in 2D, and NPCs into organoids within 12 days in 3D. Organoids contain podocytes expressing WT1, PODXL and NPHS1, distal tubules expressing the general epithelial marker EPCAM, and proximal tubules that are additionally labeled by LTL (Figure S1A-C). For KO of *WT1* we infected iPSCs harboring a Doxycyline (DOX)-inducible Cas9 protein (Ungricht et al., unpublished) with lentiviruses driving expression of a WT1-specific gRNA (gRNA1), a red fluorescent protein (RFP) and a puromycin resistance gene, and selected for viral integration in iPSCs by treating with puromycin for 6 d. Since the formation of Wilms tumor subtypes is thought to be influenced by the specific stage of kidney development in which mutations occur (Gadd *et al*, 2012), we induced genome editing by adding DOX at different stages of kidney organoid differentiation: in iPSCs prior to differentiation (KO^iPSC^), during intermediate mesoderm specification (KO^d4-7^), during NPC differentiation (KO^d9-11^) and during nephrogenesis (KO^d11-14^). *WT1* KO efficiency at the NPC stage, as determined by flow cytometry, was approximately 80% in KO^iPSC^ and 70% in KO^d4-7^ when compared to uninduced cells (Figure S1D). WT1 depletion efficiencies at d21 were comparable for all deletion time-points (Figure 1A).

**Figure 1:**
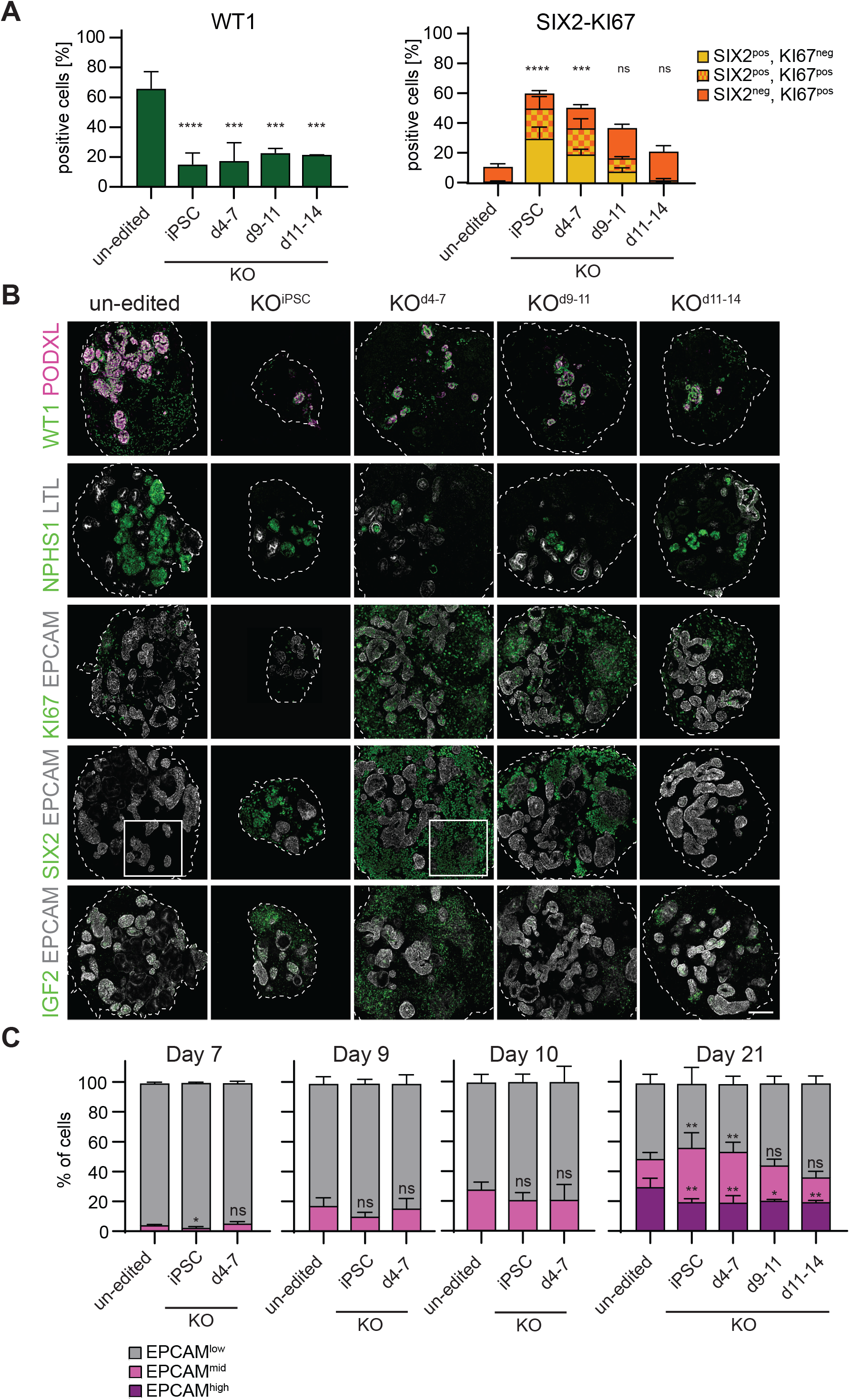
Loss of *WT1* arrests NPC progression and induces NPC overgrowth. **A:** Quantification of subpopulations in pooled d21 un-edited, KO^iPSC^, KO^d4-7^, KO^d9-11^ and KO^d11-14^ organoids by flow cytometry of indicated markers. Data is presented as mean % of positive cells +/- SD derived from n=5 (un-edited, KO^iPSC^, KO^d4-7^) or n=2 (KO^d9-11^, KO^d11-14^) independent experiments. Two-sided student’s t-test; *p*-value: ns >0.05; * ≤0.05; ** ≤0.01; *** ≤0.001; **** ≤0.0001; SIX2-Ki-67: asterisks correspond to the SIX2-KI67 population. **B:** IF staining for the indicated markers in representative un-edited, KO^iPSC^, KO^d4-7^, KO^d9-11^ and KO^d11-14^ d21 organoids. Scale bar: 100 µm. White boxed regions are shown in Figure S1J. **C:** Flow cytometry-based quantification of EpCAM^high^, EPCAM^mid^ and EpCAM^low^ populations (see gating shown in Figure S1I) in pooled organoids of indicated time-points and genotypes. Data is shown as mean % of positive cells +/- SD derived from n=3 (Day 7); n=4 (Day 9); n=2 (Day 10); n=5 (un-edited, KO^iPSC^, KO^d4-7^; Day 21) or n=2 (KO^d9-11^, KO^d11-14^; Day 21) independent differentiation experiments. Two-sided student’s t-test; *p*-value: ns > 0.05; * ≤ 0.05; ** ≤ 0.01; asterisks are placed above the respective populations (EPCAM^high^ and EPCAM^mid^).

At d21, control and *WT1* mutant organoids were of similar size, except for those derived from KO^iPSC^ cells, which were considerably smaller (Figure S1E). Culture until d30, however, revealed that KO^iPSC^ organoids remained small, that KO^d4-7^ and KO^d9-11^ organoids overgrew, and that growth of KO^d11-14^ organoids was similar to controls. Profiling with cell type-specific markers at d21 (Figure 1A,B, S1F) revealed that SIX2-positive (SIX2^pos^) cells were abundant in KO^iPSC^ and KO^d4-7^ organoids (50% and 38% of all cells, respectively), but lower in KO^d9-11^ (17%) and absent from KO^d11-14^ organoids similar to controls. In addition, the proliferation marker KI67 was elevated, in particular in KO^iPSC^, KO^d4-7^ and KO^d9-11^ organoids. Many KI67-positive (KI67^pos^) cells co-expressed SIX2. The specificity of this phenotype to depletion before d11 correlates with the downregulation of NPC genes and the induction of differentiation markers at this time-point (not shown), suggesting that overgrowth requires the deletion of *WT1* in NPCs.

PODXL- and NPHS1-expressing podocytes were strongly reduced in all *WT1* mutant organoids, including KO^d11-14^ organoids (Figure 1B). Strikingly, the few detectable PODXL-positive cells co-expressed WT1, demonstrating that they are descendants of un-edited NPCs or NPCs harboring in-frame *WT1* mutations. EPCAM^high^ and LTL-positive tubules were also reduced, indicating impaired formation of tubules (Figure 1B,C). The reduction of EPCAM^high^ cells was, however, less pronounced than the loss of podocytes. Defective nephrogenesis in KO^d11-14^ organoids in the absence of SIX2 deregulation suggests that *WT1* drives exit from the NPC state, tubular differentiation and podocyte formation through independent mechanisms. We sought to validate these findings with two independent WT1 gRNAs and found a reduction of WT1 and EPCAM^high^, maintenance of SIX2, elevation of KI67, and co-expression of SIX2 and KI67 in KO^iPSC^ organoids. Notably, phenotypic strength scaled with the KO efficiency of the respective gRNA, demonstrating on-target specificity of *WT1* KO (Figure S1G,H).

While quantifying EPCAM expression, we found a significant expansion of EPCAM^mid^ cells in KO^iPSC^ and KO^d4-7^ d21 organoids (Figure 1C, S1F,I). A significant proportion of these cells expressed SIX2 (33% and 26%, respectively). 9% of EPCAM^mid^ cells in KO^d9-11^ organoids also co-expressed SIX2, although the total number of EPCAM^mid^ cells was not significantly increased. KO^d11-14^ organoids, in contrast, were indistinguishable from controls. Immunofluorescence confirmed co-expression of EPCAM and SIX2 in cells that were arranged in epithelial-like sheets (Figure S1J), and at levels that were lower than in fully differentiated tubules. During organoid development EPCAM^mid^ cells emerged as early as d7, preceding the formation of EPCAM^high^ cells (Figure 1C). These cells are therefore likely equivalent to EPCAM^dim^ cells in the human fetal kidney (Pode-Shakked *et al*, 2017), which are kidney progenitor intermediates undergoing a mesenchymal-epithelial-transition (MET) and bridging the differentiation of EPCAM^low^ NPCs into EPCAM^high^ renal vesicle cells. WT1 is therefore required for progression beyond a pre-epithelial SIX2^pos^/EPCAM^mid^ NPC transition state, but not for initiation of MET.

While the initial induction of EPCAM was unperturbed in mutant organoids (Figure 1C), we noted increased SIX2 protein levels in NPCs at d9 (Figure S1K). It is conceivable that this SIX2 de-repression is causal for progenitor overgrowth, and that a similar mechanism underlies Wilms tumor formation in patients with neomorphic *SIX2*^Q177R^ mutations (Walz *et al*, 2015; Wegert *et al*, 2015). To test this possibility, we generated iPSCs with DOX-inducible mCherry::T2A::SIX2 and mCherry::T2A::SIX2^Q177R^ transgenes. Induction of SIX2 by adding DOX from d7, but not d9 of organoid development onwards resulted in smaller organoids that, compared to uninduced controls, had fewer EPCAM^high^- and LTL-positive tubules and WT1-positive glomeruli (Figure S2A-D). In both, d7- and d9-induced organoids, however, KI67 expression was unchanged. SIX2, despite heterogeneous and mosaic expression, was detectable in differentiated EPCAM- and LTL-positive tubule cells at d21 (Figure S2D). SIX2 overactivation is therefore not sufficient to induce overproliferation or impair NPC epithelialization.

Taken together these observations suggest that the removal of *WT1* halts NPC progression at a proliferating SIX2^high^/EPCAM^mid^ transition state, which impairs the subsequent formation of tubules and podocytes and, instead, leads to organoid hyperplasia. We found that overactivation of SIX2 was not sufficient to recapitulate these phenotypes, arguing for additional WT1 targets. Tubule and podocyte differentiation defects in KO^d11-14^ organoids indicate roles of WT1 in nephrogenesis that are independent of SIX2 silencing and consistent with WT1 stabilizing an epithelial-mesenchymal hybrid state (Sampson *et al*, 2014) and activating podocyte-specific genes (Kann *et al*, 2015).

### WT1 drives developmental transcription

We decided to define the transcriptional changes induced by absence of *WT1*, and performed RNAseq of KO^iPSC^ and KO^d4-7^ cells and respective controls at different time-points during organoid development. Principle component (PC) analysis identified PC2 associated with mesoderm specification and PC1 with further organoid development (Figure S3A). Control and KO samples were indistinguishable up to d12, but segregated along PC1 at d21. Consistent with formation of podocytes and tubules in KO organoids (Figure 1B), KO^iPSC^ and KO^d4-7^ d21 samples did not overlap with d9 NPC, d11 or d12 samples.

k-means clustering of significantly changing transcripts revealed 17 gene clusters that are dynamically regulated during organoid development and/or dysregulated in *WT1* mutants (Figure 2A,B, S3B; Table S1). These include clusters of genes that are induced or repressed during organoid development and unchanged in mutants (clusters 5, 7, 11 and 14), and clusters that are deregulated in specific cell states, in particular in d21 organoids (clusters 1,8 and 13). Notably, we did not identify clusters that are coherently mis-regulated in mutants, suggesting that WT1’s target genes depend on the developmental context. Also, expression defects in KO^iPSC^ and KO^d4-7^ organoids, despite distinct growth rates (Figure S1E), are very similar, and indistinguishable at d21.

**Figure 2:**
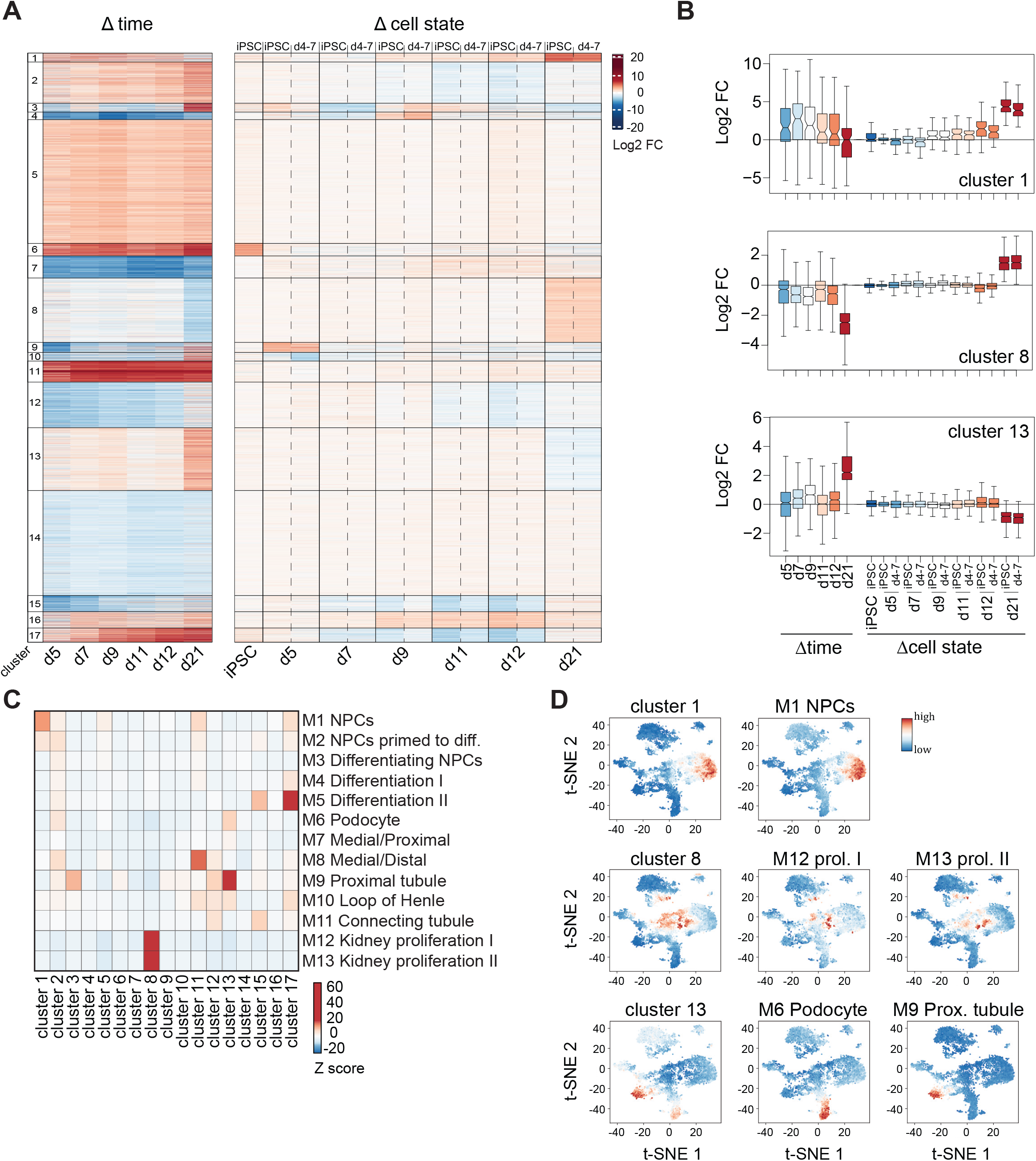
Absence of *WT1* impairs developmental transcription. **A:** k-means clustering of 7,626 genes significantly changing in any of the shown contrasts. Left: time course of wild-type organoid development; expression log2 fold changes (Log2FC) relative to iPSCs. Right: Changes in KO^iPSC^ or KO^d4-7^ cells and at indicated time-points; Log2FC relative to un-edited control at each time-point. **B:** Quantification of mRNA Log2FC of indicated gene clusters and in contrasts as specified in **A**. **C:** Gene set overlap significance scores (see Methods for details) of 17 gene clusters with fetal kidney cell type gene-sets as defined in (Lindström *et al*, 2018). Z-scores are color-coded. diff = differentiate. **D**: t-SNE maps of scRNAseq data from week 16 human fetal kidney (Hochane *et al*, 2019). Expression levels of indicated clusters and gene-sets (see **C**) are color-coded. M12 Kidney prol. I = M12 Kidney proliferation I; M13 Kidney prol. II = M13 Kidney proliferation II; Prox = proximal.

Since *WT1* KO perturbs the cellular composition of mutant organoids, we wondered if any of the gene clusters reflect cell type-specific transcription. We made use of published gene-sets that discriminate cell types of the human fetal kidney (Lindström *et al*, 2018) and calculated gene-set overlap enrichments (see Methods). This identified gene clusters 1, 8, 11, 13 and 17 to be most similar to markers of relevant cell types in the developing kidney (Figure 2C). Notably, cluster 11 was not changed, and cluster 17 only transiently deregulated in KO organoids and compensated for at d21 (Figure S3B). We validated cell type-specificity of clusters 1, 8 and 13 genes by visualizing their expression in t-distributed stochastic neighbor embedding (t-SNE) maps of single cell RNAseq (scRNAseq) of the week 16 human fetal kidney (Hochane *et al*, 2019). This confirmed co-expression with gene-sets of kidney progenitor cells (M1), proliferating intermediates (M12 and M13), and podocytes (M6) and proximal tubule cells (M9), respectively (Figure 2D).

Collectively, the transcriptional defects in developing *WT1* mutant organoids correlate with the phenotypic persistence of kidney progenitor cells (cluster 1) and reduced tubular differentiation (cluster 13). Although we cannot exclude that upregulation of cluster 8 genes reflects growth of a proliferating transit amplifying cell population, we note (1) that cluster 8 genes, similar to cell cycle-specific markers (Liu *et al*, 2017), are downregulated specifically at d21 of organoid development, suggesting cell type-independent transcription (Figure 2B, S3C), and (2) that a significant proportion of SIX2-positive cells co-express KI67 (Figure 1A), whose encoding gene *MKi-67* is a cluster 8 gene (Table S1). Upregulation of cluster 8 genes is therefore likely due to NPC hyperproliferation rather than the persistence of an additional transit amplifying cell population.

### *WT1* KO organoids recapitulate transcriptional changes in Wilms tumors

Consistent with cell type-specific expression, gene ontology (GO) term analysis (Table S1) revealed enrichment of cell cycle and metabolic processes in clusters 8 and 13, respectively (Figure S3D). Surprisingly, cluster 1 – despite containing key NPC TFs such as SIX1, SIX2, EYA1 or MEOX1 – was highly enriched for genes with skeletal muscle function, such as the TFs MYOD1 and MYOG (Table S1). Notably, expression of muscle-specific genes defines a subclass of Wilms tumors (Miyagawa *et al*, 1998; Gadd *et al*, 2012, 2017). To test if gene expression changes recapitulate Wilms tumorigenesis, we compared alterations in d21 organoids with transcriptional changes in patient tumors. For this we profiled transcription in Wilms tumor patients and respective normal tissue (WT) (Gadd *et al*, 2017). To distinguish Wilms tumors from other kidney cancer types, we also included kidney chromophobe tumors (KICH) (Davis *et al*, 2014), kidney renal papillary cell carcinoma (KIRP) (Network *et al*, 2016) and kidney renal clear cell carcinoma (KIRC) (Creighton *et al*, 2013) into the analysis. Pairwise comparison (Figure S3E) revealed that transcriptional defects in organoids correlated strongest with alterations in WT, but less with KICH, KIRP and KIRC patients. By focusing on gene clusters, we found that both the magnitude and the directionality of cluster 1 and 8 deregulation in d21 organoids was conserved specifically in WT patients (Figure 2B, 3A, S3B,F). Downregulation of the differentiation-specific cluster 13, in contrast, was also observed in KICH patient samples. Unsupervised clustering of cluster 1 genes in Wilms tumor samples (Figure 3B) identified a group of genes, including *SIX1, SIX2, EYA1* and *MEOX1*, that is strongly upregulated in all patient samples, and two groups of genes containing myogenic TFs, such as *MYOD*1 and *MYOG*, and structural muscle genes, such as *MYLPF* and *TNNC2*, that are dysregulated in only a subset of the patients.

**Figure 3:**
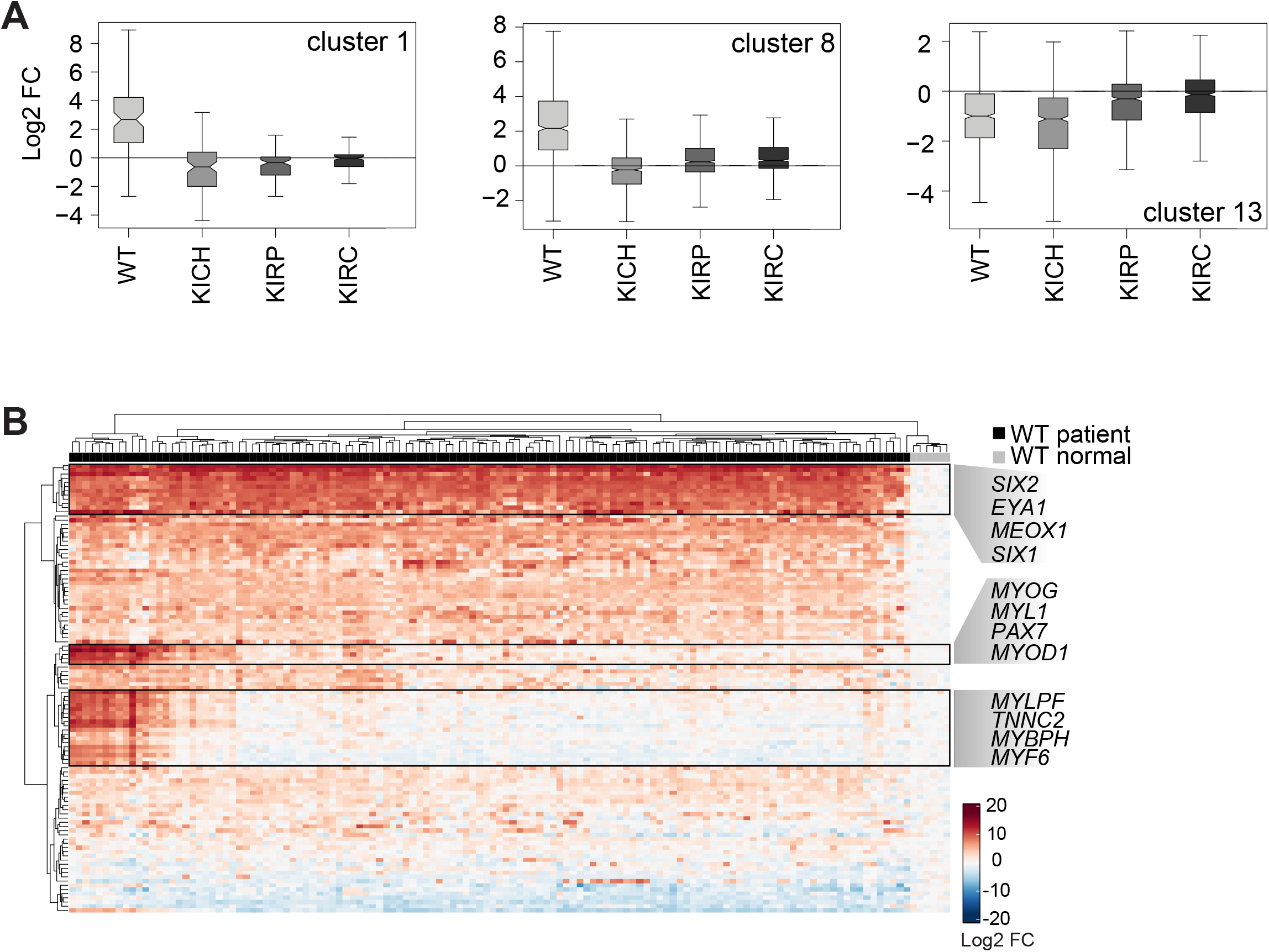
*WT1* KO organoids recapitulate the transcriptional changes of a Wilms tumor patient subgroup. **A:** Log2 FCs of indicated gene clusters in WT, KICH, KIRP and KIRC patient samples relative to the corresponding normal tissue. **B:** Unsupervised clustering of Log2FC of cluster 1 genes in WT relative to corresponding normal tissue. Clusters containing nephron progenitor genes or muscle genes are highlighted.

We therefore conclude that transcriptional changes in *WT1* mutant kidney organoids recapitulate defects in Wilms tumors, in particular the induction of NPC- and muscle-specific genes in cluster 1 and of cell-cycle-associated genes in cluster 8, and the downregulation of differentiation-specific transcripts in cluster 13.

### Niche signals propagate *WT1* mutant NPCs

The persistence of SIX2^pos^ cells and NPC transcription in *WT1* KO organoids may result from a block or a delay in developmental progression. We reasoned that serial passaging in differentiation-promoting conditions would discriminate between the two by enforcing the commitment of delayed but not of blocked mutant cells. To do so, we aggregated single cells derived from KO^d4-7^ d21 organoids and exposed them to organoid-forming conditions for 12 d (corresponding to d9-d21 of iPSC differentiation), and repeated this procedure for up to four passages (Figure 4A). Since niche signals impact tumor growth and progression (Hanahan & Coussens, 2012), we also tested the role of environment signaling in passaging of SIX2^pos^ cells. We therefore added defined ratios (0%, 10%, 25%, 75% and 90%) of GFP-expressing wild-type d9 NPCs to dissociated KO^d4-7^ organoids during the aggregation step at the beginning of each passage, expecting that the wild-type kidney structures formed by these NPCs would provide niche signals to KO cells. Quantifying the percentage of GFP-expressing cells in chimeric organoids (Figure 4B) showed stable contribution of KO cells over passages, demonstrating that KO^d4-7^ d21 organoid cells grew as fast as wild-type d9 NPCs. Un-edited control d21 organoid cells proliferated less (Figure S4A). The mutant cell type composition, in contrast, varied across passages and mixing ratios (Figure 4B,C, S4C): In the absence (0%) or presence of 10% and 25% wild-type cells, the fraction of SIX2^pos^ cells increased from approximately 30% to 60% after the first passage, but gradually declined during further passaging. Un-edited control cells, in contrast, did not gain SIX2 expression (Figure S4B). In the presence of 75% wild-type cells, the percentage of SIX2^pos^ cells remained at 30% over passages for at least 60 d. Presence of 90% wild-type cells stabilized SIX2-expression in 10% of KO^d4-7^ cells after the first passage. Therefore, paracrine signaling by wild-type cells, particularly at a wild-type:mutant cell ratio of 3:1, supports self-renewal of KO^d4-7^ SIX2^pos^ cells. At and below mixing ratios of 1:3, SIX2^pos^ cells were lost during passaging. This was not accompanied by an induction of EPCAM^high^ cells (Figure 4B) and therefore not due to overt tubular differentiation.

**Figure 4:**
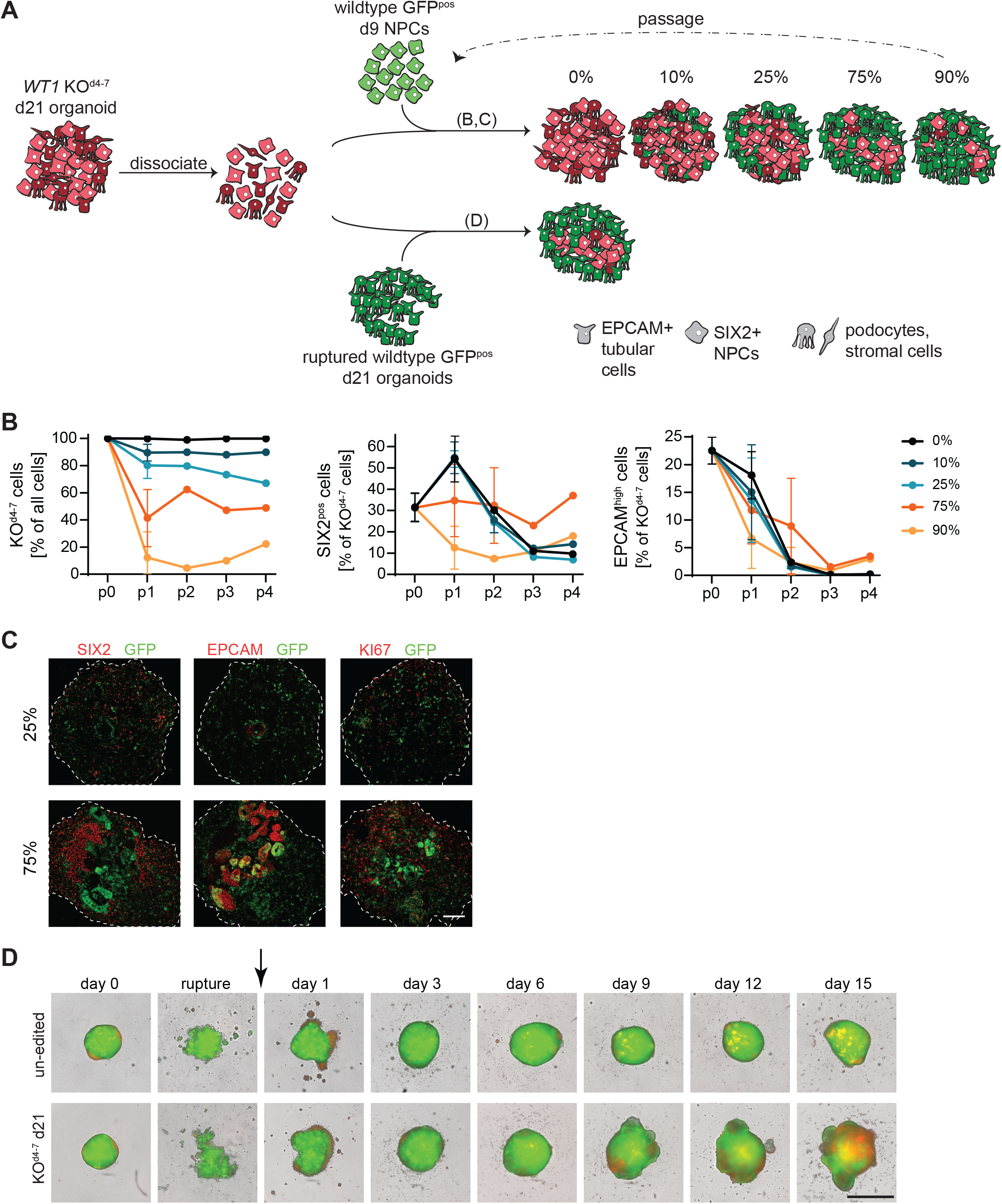
Non cell-autonomous regulation of self-renewal and heterogeneity. **A:** Schematic of the experimental flow shown in **A**-**C** (upper) and **D** (lower). **B:** Quantification of indicated cell populations in chimeric organoids after passaging with indicated ratios of wild-type GFP expressing d9 NPCs and at indicated passages. p0 indicates KO^d4-7^ d21 organoids that were used as starting material for passaging. Data is shown as mean % +/- SD from n=1 or 2 independent experiments. (% of all cells) is relative to all cells of chimeric organoids, and (% of KO^d4-7^ cells) is relative to all mutant cells, therefore excluding wild-type host cells. **C:** IF staining at p3 of indicated markers and mixing ratios as outlined in **A**. Scale bar: 100 µm. **D:** Images of organoids obtained after adding 12,500 RFP-positive un-edited d21 organoids (top) and KO^d4-7^ d21 organoids (bottom) to ruptured wild-type GFP organoids. Images were recorded at indicated time-points using an Incucyte® system. Scale bar 1mm.

Passaging in the presence of wild-type NPCs is not comparable to growth in mature tissues, such as during tumor progression or upon transplantation into PDX mouse models. We therefore decided to use ruptured GFP-expressing d21 organoids as a differentiated cell substrate for passaging of *WT1* mutant cells (Figure 4A). We estimated that d21 organoids contain 150,000-200,000 cells, to which we added 12,500 and 25,000 cells of RFP-expressing control or mutant single cells, corresponding to a mixing ratio of approximately 80-90%. KO^d4-7^ d21 cells expanded visibly (Figure 4D), expressed SIX2 and KI67 and caused organoid overgrowth (Figure 4D, S4D,E). This was in contrast to un-edited d21 cells that contributed poorly to chimeric organoids. It is possible that this difference is due to different integration efficiencies of the two cell populations and not because of unrestricted growth of mutant cells. We therefore tested control d9 NPCs and found that they integrated equally well as KO^d4-7^ d21 cells and contributed efficiently to EPCAM-positive epithelia. However, these cells did not induce organoid overgrowth or maintain SIX2 expression (Figure S4D-F). Passaging in d21 kidney organoids therefore enables persistence of SIX2 and proliferation of *WT1* mutant cells. Notably this did not require extrinsic CHIR and FGF9 (Figure S4G-I), and is therefore independent of differentiation- and growth-promoting culture conditions.

We conclude that kidney organoids can be exploited for tumor cell transplantation and growth in a developmental (d9 NPCs) and mature (d21 organoids) tissue context. Our observations identify cell autonomous and non-cell autonomous mechanisms that drive the ectopic growth of mutant NPCs, specifically a developmental block induced by absence of *WT1* and pro-self-renewal signals from a wild-type niche environment.

## DISCUSSION

KO studies in mice have identified successive functions of WT1 during nephrogenesis, while *WT1* mutations in humans predispose to familial forms of Wilms tumor and are found in 10-20% of patients (Hastie, 2017; Treger *et al*, 2019). Our *WT1*-deletion analysis in human kidney organoids consolidates previous developmental and pathological observations:

*WT1* is necessary for survival of the metanephric mesenchyme in mice (Kreidberg *et al*, 1993). KO of *WT1* in human iPSCs, in contrast, did not prevent the differentiation into NPCs (Figure S1K). Although we can’t exclude that this is because of species-specific functions of WT1 or because of non-cell autonomous rescue by unrecombined wild-type cells (Figure 1B, S1D), we propose that organoid culture conditions make up for *WT1* deficiency: Deletion of *FGF receptor 1* (*Fgfr1*) and *Fgfr2*, and of *Fgf20* and *Fgf9* in mice causes phenotypes that are reminiscent of loss of *WT1* (Barak *et al*, 2012; Poladia *et al*, 2006). WT1 directly binds to and regulates the transcription of several *Fgf* genes, and treatment of *WT1*-mutant embryonic kidneys with recombinant FGF20 rescues cell death (Motamedi *et al*, 2014). We therefore speculate that extrinsic FGF9, added between d7 and d14 of organoid formation, suppresses the pro-apoptotic effect of *WT1* loss, and exposes WT1’s oncogenic role in developmentally more advanced NPCs.

In mice, WT1 drives the epithelialization of differentiating NPCs at least in part by promoting *Wnt4* transcription (Essafi *et al*, 2011; Berry *et al*, 2015). In KO organoids, we found that a large fraction of ectopic SIX2^pos^ cells expressed EPCAM^mid^ levels and that the transcriptional induction of *WNT4* was impaired (Table S1), arguing that *WT1* deficient human NPCs are developmentally arrested while undergoing MET (Pode-Shakked *et al*, 2017). This is reminiscent of the formation of heterogeneous tumors with few epithelial elements in human patients with *WT1* mutations (Schumacher *et al*, 1997), although *WT1* lesions have been reported in epithelial cells of Wilms tumor patient-derived organoids (Calandrini *et al*, 2020). Mutant NPCs may therefore occasionally exit the progenitor state and complete MET, which is consistent with the observation that epithelialization in mutant organoids was less perturbed than glomerulogenesis (Figure 1A,C).

*WT1* deletion in mice results in hyperplasia only in the context of increased IGF2 transcription (Huang *et al*, 2016). Overgrowth of *WT1* mutant organoids, in contrast, did not require the genetic perturbation of IGF2, likely because IGF2 protein was induced in the absence of *WT1* (Figure 1B). We found that the magnitude of this induction correlated with the persistence of SIX2^pos^ cells (Figure 1B), suggesting that the two phenotypes are linked. Surprisingly, *IGF2* mRNA was not upregulated (Table S1), indicating that WT1 inhibits IGF2 expression through post-transcriptional mechanisms, such as via miRNAs (Bharathavikru *et al*, 2019).

Loss of *WT1* in humans is thought to transform immature kidney progenitor cells (Treger *et al*, 2019). The timing of *WT1* deletion, indeed, revealed that KO before d11/12 of organoid development was required for overgrowth of SIX2^pos^ cells (Figure 1A,B), matching with the terminal differentiation of endogenous NPCs. Similarly, expression of the NPC gene cluster 1 was maintained in KO^iPSC^ and KO^d4-7^ organoids from d9 onwards (Figure 2A,B). We therefore conclude that ablation of *WT1* impairs exit from the NPC state. Passaging in organoids showed that proliferating mutant SIX2^pos^ cells can be maintained long-term, but revealed the importance of environmental signaling. In particular, we were not able to propagate SIX2^pos^ cells in the absence of wild-type kidney cells. We note that primary Wilms tumor biopsies can be repeatedly passaged as spheroids (Wegert *et al*, 2020) and organoids (Calandrini *et al*, 2020) in 3D. Although growth conditions differ, it is worth considering the possibility that *WT1* KO kidney organoids recapitulate an early stage of the disease and that further tumor progression leads to independence of paracrine signals.

Although the similarity of transcriptional changes in Wilms tumor patients and *WT1* mutant kidney organoids was moderate (Figure S3E), we note that non-metanephric mesenchyme-derived cell types, such as collecting duct or vasculature, are absent from organoids (Morizane *et al*, 2015; Wu *et al*, 2018). Changes in organoids were more similar to Wilms tumors than other kidney cancers, and in particular to a subgroup of patients with upregulation of muscle-specific transcripts. Loss of *WT1* in Wilms tumor patients is associated with ectopic myogenesis (Miyagawa *et al*, 1998; Gadd *et al*, 2012), and in mouse NPCs with induction of a muscle gene expression signature (Berry *et al*, 2015). The origin of these muscle cells is unclear and may involve WT1 inhibiting mesodermal muscle differentiation (Miyagawa *et al*, 1998; Berry *et al*, 2015). Skeletal muscle progenitor TFs, such as MEOX2 and PAX7 (Chal & Pourquié, 2017), were induced only late in d21 organoids and together with muscle TFs, such as MYOG and MYOD1 (Table S1). This is inconsistent with co-development of muscle and nephrons, and suggests that muscle cells may result from the de-differentiation and ectopic re-differentiation (Shukrun *et al*, 2014) or the trans-differentiation of Wilms tumor cells.

Taken together, we here provide a framework for tracking tumor initiation and progression in human iPSC-derived organoids. Our experimental approach is restricted to pediatric cancers with few genetic inducers, limited by availability of robust organoid protocols, and confined by inductive differentiation regimes that can mask non-cell autonomous phenotypes. Tumorigenesis modeling in organoids provides access to the cancer cell type of origin, to crosstalk between different oncogenes and tumor suppressors, to interactions with the niche environment, and, eventually, to organotypic and phenotypic platforms for drug discovery and development.

## ACKNOWLEDGMENTS

Sebastien A Smallwood and team (FMI) for library preparation and sequencing, Panagiotis Papasaikas and Michael B Stadler (FMI) for help with computational analysis, Laurent Gelman (FMI) for advice and help with confocal microscopy, Melanie Rittirsch for technical assistance, Lapo Morelli (Novartis) for help with virus production and infection, Matthias Mueller (Novartis) for providing the WT29 iPSC line, and Helge Grosshans, Susan Gasser (FMI) and Soeren Lienkamp (University of Zurich) for comments on the manuscript. V.W. acknowledges support by a Boehringer Ingelheim Fonds PhD fellowship, and J.B. from the Novartis Research Foundation.

## AUTHOR CONTRIBUTIONS

VW, RU, PSH and JB conceived the study. VW designed (with input from RU) and performed all experiments and analyzed the data. RU adapted the organoid protocol and generated the WT29-iCas9 and GFP-expressing WT29-iCas9 lines. JB performed computational analyses. JB and PSH supervised the study. VW and JB wrote the manuscript with input from all authors.

## SUPPLEMENTAL INFORMATION

**Figure S1: Related to Figure 1**

**A:** Overview of the kidney organoid protocol adapted from (Morizane *et al*, 2015); Prox = proximal; Dist = distal.

**B:** IF staining of the indicated markers in representative d14, d18 and d21 organoids.

**C:** Pearson correlation coefficients of pairwise comparisons between Log2FCs of indicated organoid samples relative to corresponding iPSC samples. Expression data for d26, Morizane and d26, Takasato are from (Wu *et al*, 2018).

**D**,**G:** Quantification of WT1-expressing cells in d9 NPCs with indicated genotypes: **D** un-edited, KO^iPSC^ or KO^d4-7^ cells, and **G** KO^iPSC^ generated by indicated *WT1* gRNAs. Data presented as, **D** mean % of positive cells +/- SD from n=7 independent experiments, and **G** from one experiment.

**E:** Growth of organoids derived from un-edited, KO^iPSC^, KO^d4-7^, KO^d9-11^ and KO^d11-14^ cells between d11 and d30. Areas were derived from Incucyte® images and are presented as mean +/- SD for a minimum of n=10 organoids across all time-points.

**F**,**H:** Quantification of subpopulations expressing indicated markers in pooled d21 organoids of indicated genotypes: **F** un-edited, KO^iPSC^, KO^d4-7^, KO^d9-11^ and KO^d11-14^, and **H** KO^iPSC^ generated by indicated *WT1* gRNAs. Data is presented as, **F** mean % of positive cells +/- SD derived from n=5 (un-edited, KO^iPSC^, KO^d4-7^; KI67 and SIX2), n=2 (KO^d9-11^, KO^d11-14^; KI67 and SIX2), n=5 (un-edited, KO^iPSC^, KO^d4-7^; SIX2-EPCAM) and n=1 (KO^d9-11^, KO^d11-14^; SIX2-EPCAM) independent experiments, and **H** one experiment. Note that the SIX2-EPCAM quantification includes data presented in the Day 21 panel of Figure 1C. Two-sided student’s t-test; *p*-value: ns >0.05; * ≤0.05; ** ≤0.01; *** ≤0.001; **** ≤0.0001.

**I:** Representative flow cytometry dot plots showing EPCAM and SIX2 staining in pooled d21 organoids as in **E**. Pink boxes and associated numbers indicate % of cells that are EPCAM^high^, EPCAM^mid^ and EPCAM^low^. Blue boxes and associated numbers indicate % of cells that are SIX2^pos^/EPCAM^high^, SIX2^pos^/EPCAM^mid^ or SIX2^pos^/EPCAM^low^.

**J:** Magnification corresponding to boxes in Figure 1B. Images are overexposed to visualize EPCAM staining surrounding SIX2-positive cells (white arrowheads). Scale bar: 500 µm.

**K:** SIX2 expression in d9 NPCs by flow-cytometry. The same conditions as in **A** are shown.

**Figure S2: Related to Figure 1**.

**A:** QPCR analysis of *SIX2* transcription in d9 NPCs derived from the indicated lines that were untreated (no DOX) or treated with DOX starting from the indicated time-points. Data is presented as Log2FC relative to the untreated control. Data is shown as mean +/- SD derived from n=3 differentiation experiments.

**B:** mCherry expression in pooled d21 organoids derived the indicated lines that were untreated (no DOX) or treated with DOX starting from the indicated time-points.

**C:** Quantification of subpopulations expressing indicated markers in pooled d21 organoids derived from the indicated lines that were untreated (no DOX) or treated with DOX as in **B**. Data is shown as % of positive cells for one representative experiment.

**D:** IF staining of indicated markers in representative d21 organoids that were untreated (no DOX) or treated with DOX as in **B**. mCherry represents induction of SIX2 or SIX2^Q177R^. White arrowheads indicate expression of SIX2 in LTL^pos^ tubules. Scale bar: 100 µm.

**Figure S3: Related to Figures 2 and 3**.

**A:** PCA of un-edited (un-ed.) KO^iPSC^ and KO^d4-7^ samples at the indicated time-points.

**B**,**C:** Quantification of mRNA Log2FC of, **B** the indicated gene clusters, and **C** of cell cycle genes, and in contrasts as specified in Figure 2A. The cell cycle gene-set include G_1_S- and G_2_M-specific gene-sets (Liu *et al*, 2017).

**D:** The top six GO terms that are enriched in each of the indicated gene clusters is shown. dev. = development; proc. = process; metab. = metabolism; catab. = catabolism.

**E:** Pearson correlation coefficients of pairwise comparisons between mean Log2FCs in KO^iPSC^ and KO^d4-7^ d21 organoids relative to un-edited d21 organoids (d21 organoid), and kidney cancer patient samples relative to corresponding normal tissue. The 7’626 transcripts shown in Figure 2A were used. WT = Wilms tumor; KICH = kidney chromophobe carcinoma; KIRP = kidney papillary carcinoma; KIRC = kidney clear cell carcinoma.

**F:** Log2FCs of indicated clusters in WT, KICH, KIRP and KIRC patient samples relative to corresponding normal tissue.

**Figure S4: Related to Figure 4**.

**A**,**B:** Quantification of, **A** all, and **B** SIX2^pos^ un-edited and KO^d4-7^ cells in chimeric organoids after passaging in the presence of wild-type GFP-expressing d9 NPCs at indicated ratios after passage 1. p0 indicates KO^d4-7^ d21 organoids that were used as starting material for passaging. Data is shown as mean % +/- SD for n=2 independent experiments. **A**: Percentages are relative to all cells of chimeric organoids (% of all cells). **B**: Percentages are relative to all mutant cells (% of KO^d4-7^ cells), and relative to all un-edited control cells (% of un-edited cells). In both cases, the GFP-expressing cells are excluded from the analysis.

**C**,**D**,**I:** IF staining of the indicated markers, **C** at passage 3 and indicated mixing ratios, **D** 15 d after adding 12,500 RFP-expressing cells from un-edited d21 organoids (un-ed. d21, top), un-edited d9 NPCs (un-ed. NPCs, middle), or KO^d4-7^ d21 organoids (KO^d4-7^ d21, bottom) to ruptured wild-type GFP-expressing organoids in the presence of FGF9/CHIR, and **I** after adding 12,500 RFP-expressing cells from KO^d4-7^ d21 organoids to ruptured wild-type GFP-expressing organoids in the absence of growth factors. Scale bar: 100 µm.

**E**,**H:** Growth of organoids, **E** after adding 12,500 (12.5k) or 25,000 (25k) RFP-expressing cells in the presence of FGF9/CHIR as specified in **D**, and **H** after adding the indicated numbers of RFP-expressing KO^d4-7^ d21 organoid cells to ruptured wild-type GFP organoids and in the absence of growth factors. Areas were calculated from Incucyte® images and are presented as mean +/- standard error of the mean (SEM) for n=8 organoids per condition.

**F**,**G:** Images of organoids obtained after adding, **F** 12,500 RFP-expressing un-edited d9 NPCs to ruptured wild-type GFP organoids in the presence of FGF9/CHIR, and **G** 12,500 RFP-expressing cells from KO^d4-7^ d21 organoids to ruptured wild-type GFP organoids in the absence of growth factors. Images were recorded at the indicated time-points using an Incucyte® system. Scale bar 1mm.

**Table S1: Gene counts and log2FCs, gene clusters and GO term analysis. Related to Figure 2**.

## MATERIALS AND METHODS

### Human iPSC culture

WT29 iPSCs and their derivatives were cultured on Laminin (Biolaminin 521 LN; Biolamina #LN521) in mTeSR1 (Stem Cell Technologies; # 85850) plus 1% Penicillin-Streptomycin (Thermo Fisher #15140122). For passaging, cells were detached using TrypLE Express (Thermo Fisher #12604013), and single-cell suspensions were re-plated in mTeSR1 supplemented with 2 µM ROCK inhibitor (Y-27632 Dihydrochloride Tocris #1254). Transgenic WT29 iPSCs were generated by transfecting PiggyBac expression vector and pBase (Villegas *et al*, 2019) using Lipofectamin Stem™ (Thermo Fisher #STEM00015) in OptiMem Reduced Serum Medium (Thermo Fisher #319850629), and selected for stable integration in the presence of 100 µg/ml G418 (Thermo Fisher #10131027). Inducible cells were further selected by exposing to 1 ug/ml Doxycycline (DOX; Clonetech #631311) for 48 h and purifying the 30% of cells with mCherry expression closest to the median of the population.

### Kidney organoids

We employed a two-step protocol, consisting of an adherent (d0-d9) part forming NPCs and a suspension (d9-d21) part generating organoids, that was adapted from (Morizane *et al*, 2015) (R. Ungricht, unpublished). Induction of SIX2 and SIX2^Q177R^, and of Cas9 was achieved by treating with 1 µg/ml and 0.2 µg/ml Dox for indicated time-points, respectively.

Adherent differentiation (d0-d9): 50’000-60’000 hiPSCs/cm^2^ were plated in mTeSR1 supplemented with 2 µM ROCK inhibitor into Laminin-coated 6-well plates. After at least 6 h, medium was removed, cells gently washed with Dulbecco’s Phosphate Buffered Saline (PBS; without magnesium and calcium; Thermo Fisher #14190169), and differentiation induced by adding basic differentiation medium (BDM; advanced RPMI 1640 (Thermo Fisher #12633012), 1% Glutamax (Thermo Fisher #35050038), 1% Penicillin-Streptomycin) supplemented with 8 µM CHIR99021 (Tocris #4423) and 5 ng/ml Noggin (Peprotech #120-10C) (= d0). Medium was changed after 2 days. At d4, cells were visually inspected for the presence of contracting colonies with bright halo-like outlines, followed by medium exchange to BDM containing 10 ng/ml Activin A (R&D Systems #338_AC). At d7, medium was changed to BDM supplemented with 10 ng/ml FGF9 (R&D Systems #273-F9). At d9, corresponding to the NPC state, cells were washed, dissociated from the cell culture dish using TrypLE and counted using a Vi-CELL™ XR Cell Viability Analyzer (Beckmann).

Organoid differentiation in suspension culture (d9 onwards): 25’000 or 50’000 cells were seeded in 150 µl of BDM containing 3 µM CHIR99021, 10 ng/ml FGF9 and 2 µM ROCK inhibitor into Corning® Costar® Ultra-Low Attachment 96-well round bottom plates (Sigma #CLS7007-24EA). Surplus NPCs were frozen in CryoStor® CS10 (Stem Cell Technologies #7930). To induce aggregation, the plate was briefly centrifuged at 90 g for 3 minutes (min). At d10 (d1 in suspension), 100 µl of medium was replaced with 150 µl of BDM plus 10 ng/ml FGF9. At d11 (d2), 100 µl of medium was replaced with 100 µl of BDM plus 10 ng/ml FGF9. At d14 (d5), d16 (d7) and d18 (d9), 100 µl of medium was replaced with 100 µl of BDM without growth factors. During culture beyond d21, medium was changed 3 times per week. Organoid growth was determined using an Incucyte® system: Brightfield images were recorded every 24h, or as indicated, and organoid sizes were calculated. Object recognition parameters were manually defined for each experiment, and the detected objects validated.

To generate single cell suspensions for flow cytometry or passaging, organoids were transferred into tubes using a cut P-1000 tip, and washed twice with PBS. A 1:1 mix of non-enzymatic cell dissociation solution (Thermo Fisher #13151014) and 0.25% Trypsin-EDTA (Thermo Fisher #25200056) was added for 10 mins at 37°C, and organoids were dissociated by pipetting up and down ten times. Trypsin was inactivated by adding 10% fetal calf serum (FCS; Bioconcept #2-01F36-I) in PBS, washed with 1% FCS and passed through a 50 µm filter (BD Biosciences #340632).

### Chimeric organoids

For mixing with NPCs (Figure 4B,C, S4A-C), single cell suspensions of RFP-expressing un-edited or KO^d4-7^ d21 organoids were generated as described above, aggregated with freshly thawed GFP-expressing WT29-iCas9 d9 NPCs at indicated ratios to a total of 50,000 cells, and plated into Ultra-Low Attachment 96-well round bottom plates in 150 µl BDM, supplemented with 3 µM CHIR99021, 10 ng/ml FGF9 and 2 µM ROCK inhibitor. Culture was continued as detailed in «Organoid differentiation in suspension culture» above. At d12 of suspension culture, organoids were dissociated into single cell suspensions that were used for flow cytometry analysis and for passaging by aggregating with freshly thawed GFP-expressing WT29-iCas9 d9 NPCs at respective ratios and subjecting to suspension culture. This was repeated for up to four passages. After each passage, a minimum of six organoids was processed for cryosectioning and immunofluorescence staining, as described below.

For mixing into d21 organoids (Figure 4D, S4E-I), GFP-expressing WT29-iCas9 d21 were mechanically ruptured by pipetting them up and down five times in a P-200 tip. RFP-expressing un-edited and KO^d4-7^ d21 organoids were dissociated into single cell suspensions as described above, and un-edited RFP-expressing d9 NPCs were freshly thawed. After cell counting, indicated cell numbers were added to ruptured organoids, plates briefly spun to induce aggregation, and culture in BDM resumed as indicated, either in the presence or absence of 3µM CHIR99021 and 10ng/ml FGF9.

### Molecular biology

The coding sequence of SIX2 was amplified from human iPSC cDNA, T2A sequences and Gateway cloning sites added by polymerase chain reaction (PCR), and recombined into pDONR221 using Gateway technology (Thermo Fisher #11789020 and #11791020). The Q177R point mutation in SIX2 was introduced by PCR. Expression vectors were generated by recombining with a PiggyBac pPB-TRE-mCherry-DEST-rTA-HSV-neo expression destination vector. *WT1* gRNA-encoding vectors are derived from pRSI16-U6-sh-UbiC-TagRFP-2A-Puro (Cellecta #SVSHU616-L).

Oligonucleotide sequences:

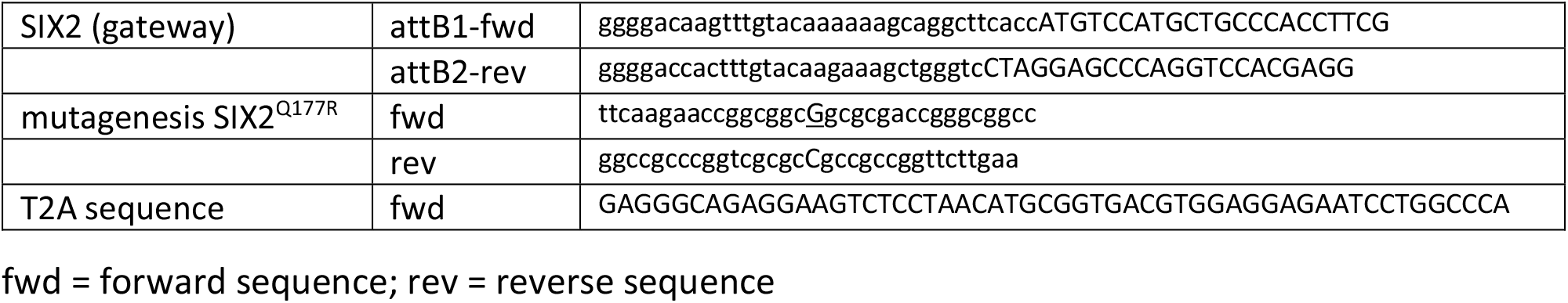

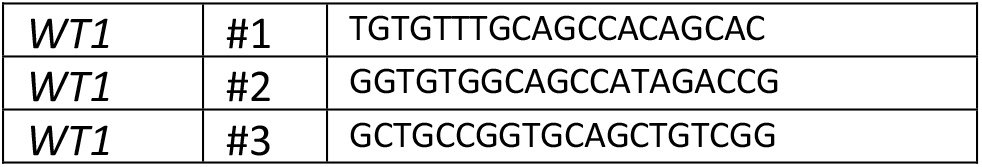

### Lentivirus production and human iPSC *transduction*

Lentiviruses were produced in HEK293T cells. Prior to transfection, HEK293T cells were seeded onto collagen I-coated 6-well tissue culture plates (BD biosciences #346400) in packaging medium (DMEM (Thermo Fisher #11965), supplemented with 10% FCS and 1% Non-essential amino acids (Thermo Fisher #11140050)). The next day, cells were transfected with *WT1* gRNA-encoding vectors and Cellecta packaging mix (Cellecta #CPC-K2A) using the TransIT™293 transfection reagent (Mirus Bio #MIR 2700) in OptiMem Reduced Serum Medium (Thermo Fisher #31985062). The next day, medium was changed to 1 ml of packaging medium. After 3 days (d), the virus-containing supernatant was collected, filtered through a 50 µm filter and stored at -80°C.

For virus titration, WT29-iCas9 cells were seeded into 6-well plates (200,000 cells / well) in mTeSR1 medium supplemented with ROCK inhibitor. After 7 h, different volumes of viral supernatant were added to the cells. After 3 d with daily mTeSR1 medium changes, cells were detached and RFP fluorescence was measured by flow cytometry. Based on this titration, WT29-iCas9 cells were transduced at a multiplicity of infection of 0.5, and infected cells were selected with puromycin (Thermo Fisher #A11138-03) for 6 days.

### RNA isolation, cDNA synthesis, qPCR

RNA was isolated using RNeasy Mini Kits (Qiagen #74104) and RNase-Free-DNase Sets (Qiagen #79256) according to the manufacturer’s instructions, and concentration determined using a NanoDrop (Thermo Fisher).

cDNA was generated from at least 400 ng of total RNA using SuperScript III Reverse Transcriptase (Thermo Fisher # 18080044) using oligodT priming. 25 ng of cDNA was subjected to quantitative PCR (qPCR) on a Step One Plus™ Real-Time PCR System (Thermo Fisher) using the TaqMan Fast Universal PCR Master Mix (Thermo Fisher # 4364103). Expression of SIX2 was quantified in technical duplicates using Universal Probe Library (UPL, Roche) Probe 88 together with custom-designed primer pairs: Fwd: ggcaagtcggtgttaggc, Rev: ggctggatgatgagtggtct, and multiplexing with a *GAPDH* probe (Thermo Fisher # 4326317E). Cycle threshold (CT) values were normalized to *GAPDH* and to controls (ΔΔCT method).

### Flow cytometry

Single live cell suspensions were washed with 1% FCS in PBS and resuspended in flow cytometry buffer (2% FCS and 1 mM ethylenediaminetetraacetic acid (EDTA; Thermo Fisher #AM9260G) in PBS).

For flow cytometry analysis of stained cells, dissociated cells were fixed using the BD Cytofix/Cytoperm Fixation/Permeabilization Kit (BD Biosciences #554714) according to the manufacturer’s instructions with the exception that 0.4% bovine albumin fraction V solution (BSA, 7.5%, Thermo Fisher #15260037) in PBS was added to fixed cells before centrifugation. Cells were incubated with primary antibodies in permeabilization buffer at 4°C for 60 mins, washed three times, incubated with secondary antibodies at 4°C for 60 mins, washed three times and resuspended in permeabilization buffer for flow analysis. Antibodies were EPCAM-Alexa Fluor (AF) 647 (Abcam #ab239273, 1:200); Ki-67-FITC (eBisoscience #11-5698-82, 1:200); SIX2 (Proteintech #11562-1-AP, 1:100); WT1 (Abcam #ab89901, 1:200); donkey-anti-rabbit AF488 (Thermo Fisher #A-21206, 1:500); donkey-anti-rabbit AF 594 (Thermo Fisher #A-21207, 1:500); donkey-anti-rabbit AF 647 (Thermo Fisher #A-31573, 1:500). Analytic flow cytometry was performed on a BD LSRFortessa™ or BD LRSII™ device, cell sorting on a BD FACSAria™ Fusion Cell Sorter. Data was analyzed in Flow Jo.

RFP-expression in fixed *WT1* KO cells of chimeric organoids (Figure 4B, S4A,B) by flow cytometry was ambiguous. We therefore quantified the fraction of mutant cells by determining the percentage of GFP-expressing wild-type WT29-iCas9 cells and correcting for heterogeneous GFP expression.

### Immunofluorescence

Kidney organoids were pooled, washed twice with PBS, and fixed with 4% paraformaldehyde (PFA) in PBS for 20 mins at 4°C. After washing three times with PBS, PBS was removed, organoids were resuspended in 50% sucrose (Sigma #84097) in PBS and stored at 4°C overnight. The next day, organoids were embedded in gelatin solution (7.5% gelatin from porcine skin (Millipore #48722) and 10% sucrose in PBS) overnight at 4°C. The next day, organoids were mounted with Q Path Tissue OCT Medium (VWR #0011243) to generate frozen blocks which were cut into 10-14 µm sections using a Leica CM3050S cryostat. The sections were washed with PBS for 10 mins at room temperature (RT). When using biotinylated primary antibodies, slides were incubated with blocking/permeabilization buffer (1% BSA and 0.2% Triton X-100 in PBS) for 15 mins, then with blocking/permeabilization buffer containing four drops per ml of Streptavidin Block solution (Streptavidin/Biotin Blocking Kit; Vectorlabs #SP-2002) for 15 mins, and then with blocking/permeabilization buffer containing four drops per ml of Biotin Block solution for 15 mins. When using un-biotinylated primary antibodies, slides were instead incubated with blocking/permeabilization buffer for 30 mins at RT. After a quick wash in PBS slides were incubated with primary antibodies in 1% BSA in PBS for 1 h at RT. Afterwards, slides were washed twice with PBS for each 10 mins, incubated with secondary antibodies and Hoechst 33342 diluted into 1% BSA in PBS for 1 h at RT, and after two additional washes with PBS for 10 mins each, mounted in ProLong™ Diamond Antifade Mountant (Thermo Fisher #P36970).

Antibodies were: CDH1 (BD Biosciences #610181, 1:200); EPCAM-AF647 (Abcam #ab239273, 1:200); Hoechst 33342 (Thermo Fisher #H3570; 1:10000); HOXD11 (Sigma #SAB1403944; 1:300); IGF2 (Thermo Fisher #MA5-17096; 1:200); Ki-67 (SolA15, eBisoscience #14-5698-82, 1:200) (Ki-67 / Ki-67-FITC (SolA15, eBisoscience #14-5698-82, 1:200); LHX1 (OriGene #TA504528; 1:1000); LTL-Biotinylated (Vectorlabs B-1325, 1:500); LIN28A (Cell Signaling #3978, 1:600); LIN28B (Cell Signaling #4196, 1:250); NPHS1 (R&D Systems #AF4269, 1:60); PAX2 (Invitrogen #71-6000; 1:100); PAX8 (Proteintech #10336-1-AP; 1:100); PODXL (R&D Systems #AF1658, 1:500); SIX1 (Cell Signaling #12891; 1:200), SIX2 (Proteintech #11562-1-AP, 1:100); WT1 (Abcam #ab89901; 1:200); donkey-anti-rabbit AF 488 (Thermo Fisher #A-21206, 1:500); donkey-anti-rabbit AF 594 (Thermo Fisher #A-21207, 1:500); donkey-anti-rabbit AF 647 (Thermo Fisher #A-31573, 1:500); donkey-anti-goat AF 594 (Thermo Fisher #A-11058, 1:500); donkey-anti-goat AF 647 (Thermo Fisher #A-21447, 1:500); donkey-anti-sheep AF 488 (Thermo Fisher # A-11015, 1:500); donkey-anti-mouse AF 488 (Thermo Fisher #A-21202, 1:500); donkey-anti-mouse AF 594 (Thermo Fisher #A-21203, 1:500); donkey-anti-mouse AF 647 (Thermo Fisher #A-31571, 1:500); donkey-anti-rat AF 647 (Abcam #150155, 1:500); Streptavidin Fluorescent Dye 633-I (Abnova #U0295, 1:500). Images were acquired on a Zeiss LSM710 scanning head confocal microscope and handled with Fiji software.

### Bioinformatics

RNA isolation of three independent biological replicates was performed as described above and RNA-seq libraries prepared using the TruSeq mRNA Library preparation kit (Illumina #20020595). RNA sequencing was performed on an Illumina HiSeq2500 machine (50 bp single-end reads). RNA-seq reads were aligned to the human hg38 genome using *qAlign* from the Bioconductor package QuasR (Gaidatzis *et al*, 2015) with default parameters except for *aligner=“Rhisat2”* and *splicedAlignment=TRUE*. Alignments were quantified for known UCSC genes obtained from the TxDb.Hsapiens.UCSC.hg38.knownGene package using *qCount* (Table S1).

Principle component analysis (PCA) (Figure S4A) using normalized read counts of merged replicates and considering the top 30% variable genes was performed using the prcomp function in R.

Differential gene expression was determined using edgeR (Robinson & Oshlack, 2010). For heatmap visualization (Figure 2A), 7’626 genes were considered that were significantly regulated during control organoid formation (absolute log2 fold gene expression change (log2FC) at d5, d7, d9, d11, d12 or d21 relative to iPSCs greater than log2(3) with a false discovery rate (FDR) smaller than 0.001) or upon *WT1* KO (absolute log2FC in KO^iPSC^ or KO^d4-7^ organoids relative to control organoids at any time-point greater than log2(3) with a FDR smaller than 0.001).

For comparison with published kidney organoid gene expression (Wu *et al*, 2018) (GSE118184) (Figure S1C), scRNAseq reads of 218 iPSCs and of 25120 (Morizane protocol) and 82024 (Takasato protocol) cells from d26 organoids were normalized and collapsed, and log2FCs relative to iPSCs were calculated using a pseudocount of 1. Correlation coefficients are based on 12817 genes detected in (Wu *et al*, 2018) and the RNA-seq dataset reported in this work. Pearson correlation coefficients were calculated using R’s *cor* function

For generation of gene-set overlap scores (Figure 2C), we first calculated the odds-ratio for each pairwise comparison between embryonic kidney gene sets (Lindström *et al*, 2018) and gene clusters 1-17, using the *fisher*.*test* function in R. To correct for biases introduced by different gene-set sizes, each observed odd-ratio was then normalized by calculating a Z-score: z_score = (obs – mean_rand) / sd_rand, where obs is the observed odds-ratio for a given pairwise comparison, and mean_rand and sd_rand are the mean and standard-deviation of 100 randomized odd-ratios, obtained from equal-sized sets of randomly selected genes.

Analyses of enriched gene sets (Figure S3D and Table S1) was performed using DAVID (Huang *et al*, 2008) and selecting GOTERM_BP_ALL.

Single-cell RNAseq (scRNAseq) datasets of week 16 fetal kidney scRNAseq (Hochane *et al*, 2019) (GSE114530) were integrated with and contrasted to the results of this study. For the analysis and visualization of the scRNAseq data (Figures 2D) we filtered out genes detected in < 1% of the cells as well as the abundantly expressed and noisy ribosomal protein genes. From the remaining fraction, only the top 5% overdispersed genes were selected as input for the downstream dimensionality reduction and dataset integration methods according to a mean-variance trend fit using a semi-parametric approach (Zheng *et al*, 2017).The coordinates of the first 32-principal components were used to obtain the 2D tSNE representation of the data. For visualization purposes per-cell gene expression values were subjected to k-nearest neighbor smoothing (k=64) and normalized against a random set of 2000 detected genes in order to control for artefactual expression gradients.

For transcriptional changes in kidney cancer patients (Figure 3, S3E,F), Wilms tumor (WT; Gadd et al., 2017), Kidney Chromophobe Carcinoma (KIRC; TGCA; Davis et al., 2014), Kidney Papillary Cell Carcinoma (KIRP; The Cancer Genome Atlas Research Network, 2016), and Kidney Clear Cell Carcinoma (KIRC; TCGA; Creighton et al., 2013) datasets (TARGET-WT, TCGA-KICH, TCGA-KIRP, and TCGA-KIRC) were downloaded from GDC (https://portal.gdc.cancer.gov) using the TCGAbiolinks package available from Bioconductor. Data sets were normalized, and log2FCs for each sample calculated over the mean of the respective control samples using a pseudocount of 1. Correlation coefficients are for the 7’626 genes defined in Fig. 3A, and the mean of log2FCs in KO^iPSC^ and KO^d4-7^ d21 organoids. Unsupervised clustering was performed using the Heatmap function from the Bioconductor package ComplexHeatmap.

## DATA AVAILABILITY

The RNA-seq data reported in this study have been deposited at ArrayExpress with the following accession number: E-MTAB-9957, and can be accessed with the Username: Reviewer_E-MTAB-9957, and the Password: Jrpdwjbn.

