## Supplementary figures and images for "Wilms Tumorigenesis in Human Kidney Organoids"

### Figure S1

#### Waehle et al., Supplemental Figure 1

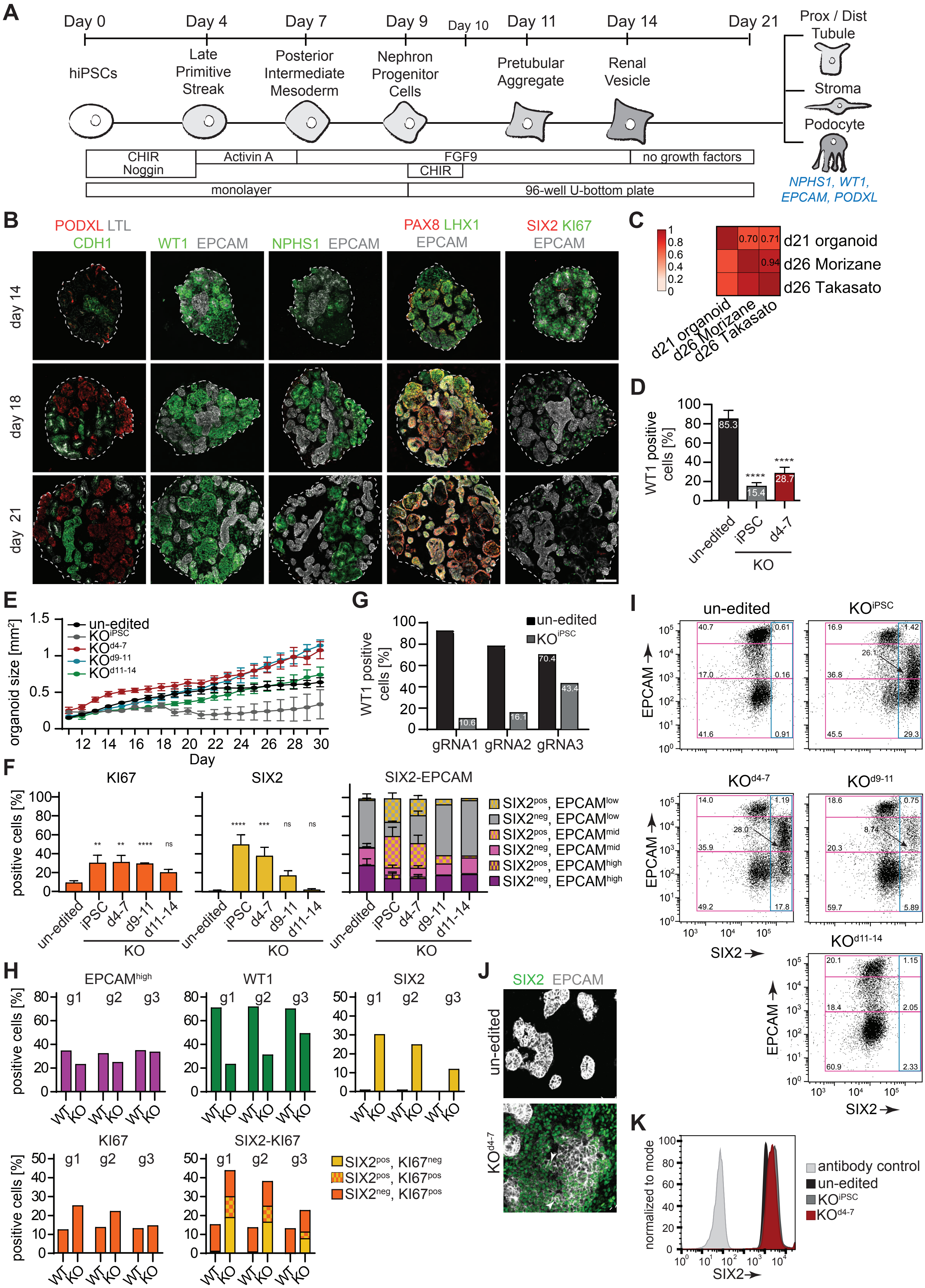

### Figure S2

#### Waehle et al., Supplemental Figure 2

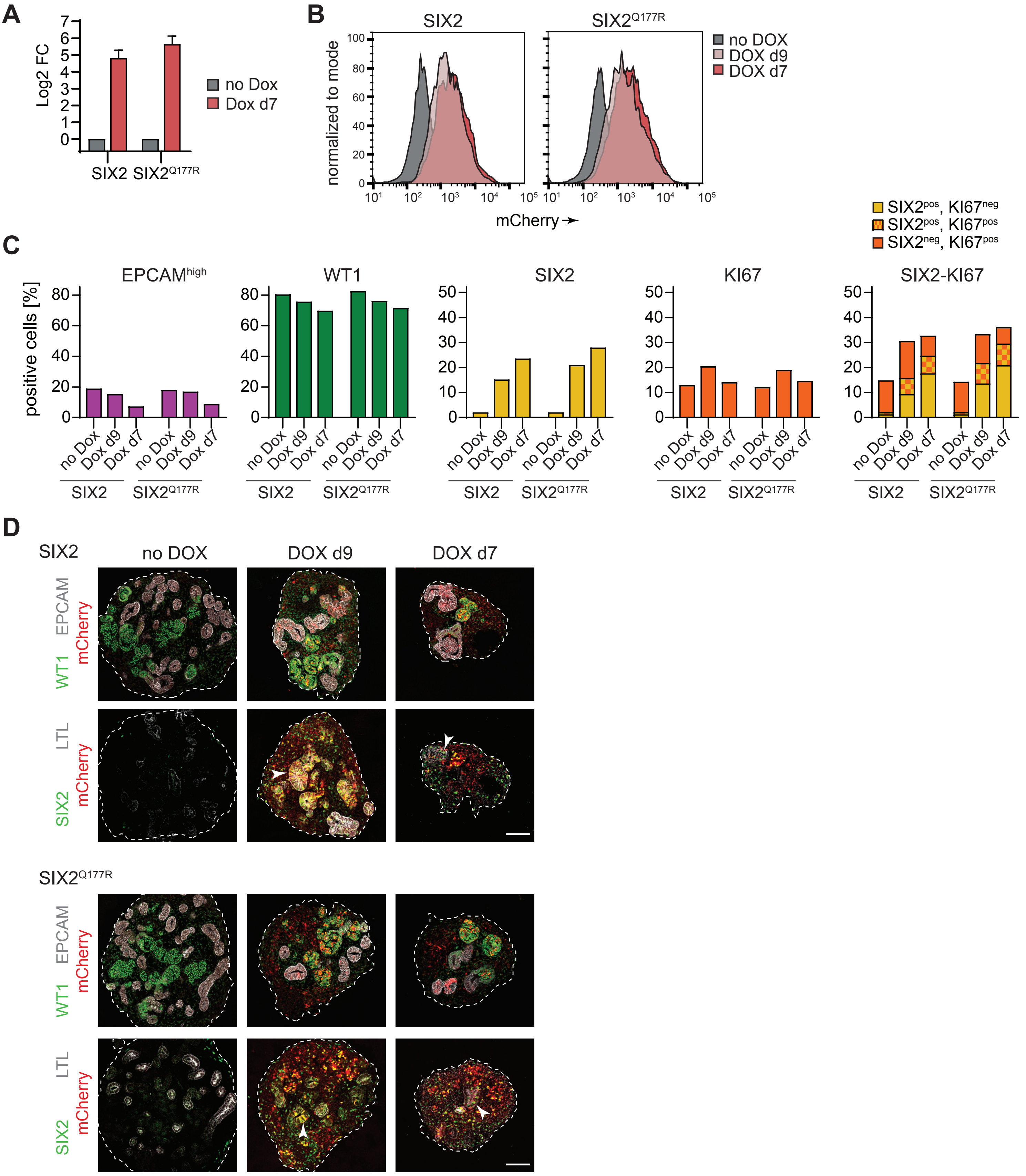

### Figure S3

#### Waehle et al., Supplemental Figure 3

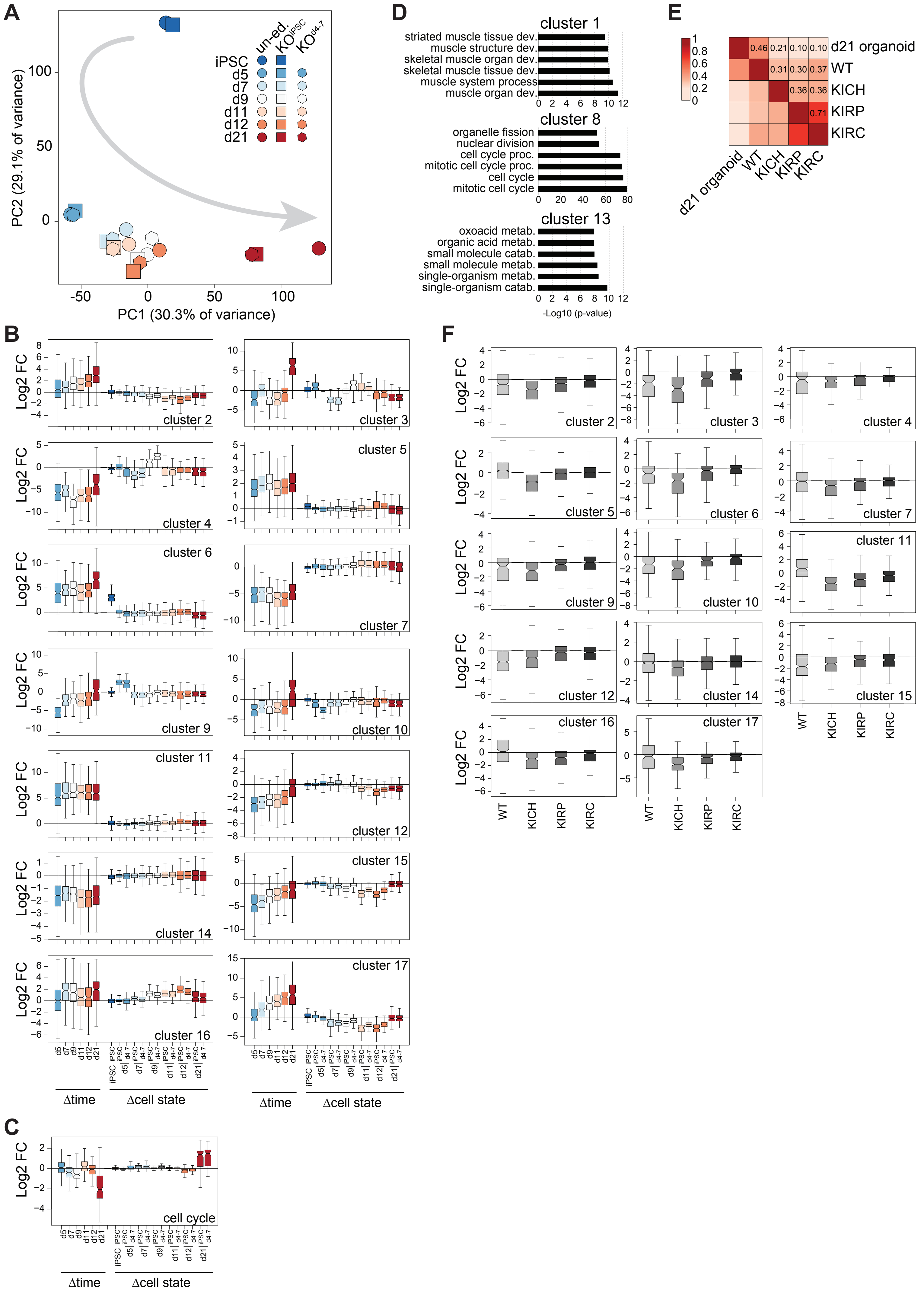

### Figure S4

**Waehle et al., Supplemental Figure 4**

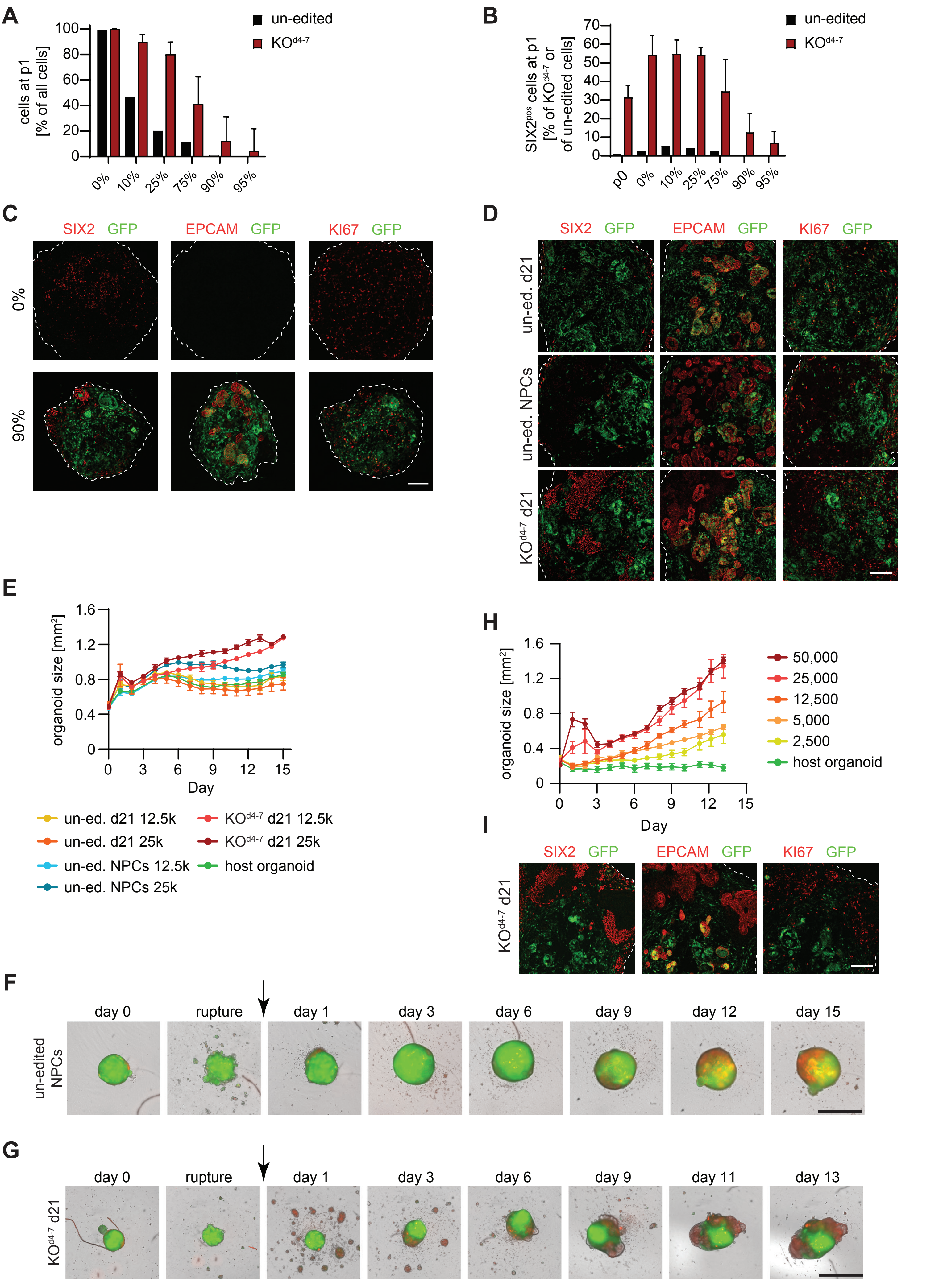
